# Food Restriction Augmented Alpha1–Adrenergic Mediated Contraction in Mesenteric Arteries

**DOI:** 10.1101/2021.12.12.472277

**Authors:** Rany Vorn, Hae Young Yoo

## Abstract

Food restriction (FR) enhances the sensitivity to cardiopulmonary reflexes and α1- adrenoreceptors in the female, despite hypotension. The effect of male FR on cardiopulmonary and systemic vascular function is not well understood. This study examines the effects of FR on cardiopulmonary, isolated mesenteric arterial function and potential underlying mechanisms. We hypothesized that FR decreased eNOS activity in mesenteric arteries. Male Sprague Dawley (SD) rats were randomly divided into three groups: (1) control (n=30), (2) 20 percent of food reduction (FR20, n=30), and (3) 40 percent of food reduction (FR40, n=30) for five weeks. Non-invasive blood pressure was measured twice a week. Pulmonary arterial pressure (PAP) was measured using isolated/perfused lungs in rats. The isolated vascular reactivity was assessed in double-wire myograph. After five weeks, food restricted rats exhibited a lower mean arterial pressure and heart rate, however, only FR40 groups exhibited statistically significant differences. The basal tone of PAP and various vasoconstrictors did not show significant differences in pulmonary circulation between each group. We observed that food restriction were enhanced the sensitivity (EC_50_) in response to α1-adrenoreceptors (phenylephrine, PhE)-induced vasoconstriction, but not to serotonin, U46619, and high K^+^ in the mesenteric arteries. FR reduced endothelium-dependent relaxation via decreased function of endothelial nitric oxide synthase (eNOS)-nitric oxide (NO) pathway in the mesenteric arteries. PhE-mediated vasoconstriction in mesenteric arteries was eliminated in the presence of eNOS inhibitor (L-NAME). In addition, incubation with NOX2/4 inhibitors (apocynin, GKT137831, VAS2870) and reactive oxygen species (ROS) scavenger inhibitor (Tiron) were eliminated the differences of PhE-mediated vasoconstriction but not to cyclooxygenase inhibitor (indomethacin) in the mesenteric artery. Augmentation of α1–adrenergic mediated contraction via inhibition of eNOS-NO pathway by increased activation of ROS through NOX2/4 in response to FR. Reduced eNOS-NO signaling might be a pathophysiological counterbalance to prevent hypovolemic shock in response to FR.

## INTRODUCTION

Cardiovascular disease (CVD) remains a major problem worldwide [1], and in some studies, is reported to be caused by vascular endothelial and smooth muscle cells dysfunction [2, 3]. Endothelial cell dysfunction plays a key role in regulating vascular reactivity [4]. Vascular endothelium also has a significant role in regulating vascular tone by releasing a balanced production of vasoconstrictor and vasodilator factors [5]. eNOS is highly expressed in the vascular endothelium to maintain vascular dilation and protect against vascular disease [6, 7]. In pathological conditions, dysfunction of eNOS enzyme leads to oxidative stress [8], which increases production of reactive oxygen species (ROS) induced nitric oxide (NO) bioavailability in endothelial cells [9].

Nicotinamide adenine dinucleotide phosphate oxidases (NOXs) is a major source of ROS, including superoxide and hydrogen peroxide production in the endothelial cells [10]. Seven isoforms of NOXs are identified; only NOX1, NOX2, NOX4, and NOX5 are expressed in vascular endothelial cells [11, 12]. NOXs may be involved in regulating vascular reactivity by upregulation of ROS production in pathological conditions such as hypertension and diabetes [13-15]. At the physiological level, ROS production is important for maintaining vascular endothelial cells and smooth muscle cells homeostasis [16].

Food restriction (FR) is a reduction of daily food intake, and may maintain either the normal micronutrient status, or may induced malnutrition. FR has been widely used in nutritional studies of anorexia nervosa (AN) [17] and non-pharmacological treatment in various diseases such as hypertension [18], obesity [19], and diabetes [20]. FR is thought to lengthen the lifespan in animal studies by 20–40% [21, 22]. Clinical and animal studies suggest that FR is associated with reducing atherosclerosis risk factors by increasing high-density lipoprotein cholesterol and decreasing low-density lipoprotein cholesterol, triglycerides and lowering blood pressure [23, 24].

Despite FR having beneficial effects on various diseases [18, 20, 25], adverse effects of FR on the cardiovascular system have also been reported [26, 27]. Abnormal cardiac function and structure have been observed in both animal and human malnutrition [28, 29]. FR contributes to the development of cardiovascular events such as hypotension [26, 30], and myocardial dysfunction [27]. FR induces lower production of NO in the blood plasma and increases cardiopulmonary reflexes in response to α1–adrenergic activity in female rats [24]. A recent study suggested that enhanced α1–adrenergic receptor might be a counterbalance to maintaining blood pressure in response to FR induced hypotension [24]. The α1-adrenergic receptor is a neurotransmitter which plays an important role in regulation of blood flow and essentially in controlling blood pressure [31, 32]. Few studied have investigated the pulmonary arterial pressure and vascular reactivity of systemic arteries such as the mesenteric artery in male FR. This study investigated the effects of FR from 20% to 40% on pulmonary vascular function and isolated mesenteric arteries. The influence of FR on whole organ blood flow and direct evaluation of VSMCs and endothelial cells without the influence of surrounding adipose tissue in the mesenteric arteries is not well understood. We hypothesized that FR attenuated endothelium-dependent relaxation and enhanced mesenteric vascular contractility. We also tested the effects of eNOS, NOXs, and ROS inhibitors in response to FR, in order to understand the physiological changes in vascular contractility.

## MATERIALS AND METHODS

### Animal Preparation

Male Sprague-Dawley (SD) rats were purchased at 7 weeks of age. After one week environmental adjustment, 90 SD rats were randomly divided into 3 groups: 1) control (the rats were allowed unrestricted access to tap water and food); 2) FR20 (20 percent of food restriction) ; and 3) FR40 (40 percent of food restriction). The food servings were calculated based on daily monitoring of food intake during the first week, and were adjusted in the remaining weeks of the study. FR20 groups received an average of 18.4 g of food per day and FR40 groups received an average of 13.8 g of food per day for 5 weeks with unrestricted access to water. The animals were housed individually to assure food intake during a 12-h light/12-h dark cycle.

### Non-Invasive Blood Pressure Measurement

In this experiment, weekly blood pressure was monitored by using a non-invasive blood pressure system the tail cuff method (CODA, Kent scientific Corp., USA). Rats were allowed to adjustto the plastic restrainer for a week before the experiment. After adapting to the plastic restrainer, rats were measured at 32 °C–35 °C after stabilization.

### Pulmonary Arterial Pressure Measurement

The rats were anesthetized with an intraperitoneal injection of 90 mg/kg of ketamine with 10mg/kg of xylazine before the experiment. The animal was confirmed to be fully anesthetized by limb withdrawal using toe pinching. After the animals were fully anesthetized, a tracheotomy was performed, and animals were ventilated with an inspiration gas mixture (21 % O_2_, 5% CO_2,_ and N_2_ balanced) via a rodent ventilator (Respirator 645, Harvard Apparatus, USA). The rats were continuously maintained at 85 breaths/min and tidal volume at 10 ml/kg. Sternotomy was performed and heparin (200 U/kg) was quickly injected into the right ventricle to prevent blood coagulation. After injection with heparin, a cannula was quickly inserted into the pulmonary artery and tightened with the surgery silk. An additional cannula was inserted immediately into the left atrium via left ventriculotomy to maintain the constant perfusion of blood flow. The perfusate composed 30 ml of physiological salt solution (PSS) and 20 ml of whole blood was perfused via a peristaltic pump (Servo amplifier 2990, Harvard Apparatus, USA). The flow rate of the perfusate was set at 15 ml/min at rest. The mean pulmonary arterial pressure (PAP) was measured continuously using a pressure transducer (Abbott pressure transducer, Abbott Lab, USA) and the data acquisition and storage was done with Powerlab/4ST and Chart 5 (AD Instruments, Australia). After achieving a stable basal PAP, a normoxia and repetitive hypoxia-induced pulmonary vasoconstriction (HPV) was applied with a hypoxic gas mixture (3% O_2_, 5% CO_2_, and 92% N_2_) in 5 min intervals. As a preconstrictor, angiotensin II (Ang II, 1 µg) was applied, and the perfusate temperature was maintained at 37°C. The PSS used for ventilated and perfused lung consisted of the following (in mM/L): NaCl 120, KCl 4.3, MgSO_4_ 1.0, KH_2_PO_4_ 1.1, NaHCO_3_ 19, Glucose 10, MgCl_2_ 1.2 and CaCl_2_ 1.8.

### Vascular Tissue Preparation

The intestine was quickly removed and placed in Normal Tyrode (NT) solution at 4 °C. The second and third branches of the mesenteric arteries were quickly isolated and cut into segments of 2.5∼3 mm in length (inner diameter: 200∼250 μm). The composition of NT solution contained the following: (composition in mM/L: NaCl 140, KCl 5.4, NaH_2_PO_4_ 0.33, 4-(2-hydroxyethyl)-1-piperazineethanesulfonic acid (HEPES) 10, Glucose 10, MgCl_2_ 1.0 and CaCl_2_ 1.8 (adjusted with NaOH to pH 7.4).

### Vascular Function Measurement

The vascular function measured in resistance artery using the Multiwire Myograph System (DMT 620 M; Aarhus, Denmark). The arterial rings were mounted on two 25 μm tungsten wires with 5 ml of PSS in each chamber. Arteries were equilibrated at an intraluminal pressure of 50 mmHg to 70 mmHg (0.5–0.7 g) using a micromanipulator. A segment of arterial rings were stabilized in PSS with continuous gas bubbling (21 % O_2_. 5% CO_2_ and N_2_ balanced at 37 °C) to maintain pH at 7.4. The composition of PSS solution contained the following (composition in mM/L: NaCl 118, KCl 4, MgSO_4_ 1.0, NaH_2_PO_4_ 0.44, NaHCO_3_ 24, Glucose 5.6, and CaCl_2_ 1.8.). The isometric tension was recorded by a data acquisition system (Lab Chart Pro version 8.0). The arterial rings were tested for the validity of segment preparation by determining response to a high concentration of potassium chloride (80 mM K^+^-PSS). A high concentration of 80 mM K^+^ was prepared by replacing NaCl with equimolar KCl. The rings were washed with PSS solution until resting tone returned. The endothelial integrity was observed with a single dose of phenylephrine (PhE, 10 μM) and acetylcholine (ACh, 10 μM). Endothelium-dependent relaxation was then assessed in all vessels by response to ACh (0.05–1 μM) that were precontracted 10 μM of phenylephrine (PhE). Endothelium-independent relaxation was assessed with sodium nitroprusside (SNP, 0.05–10 μM). To assess the vascular contractility, the vasoconstrictors were applied in each vessel, such as PhE (0.05–10 μM), thromboxane A2 analogue (U46619, 1 nM–1 μM), 5-hydroxytryptamine (5-HT, 0.05–10 μM) and high potassium chloride (KCl, 20–120 mM). In some experiment, arteries were preincubated for 30 minutes with 100 μM of nonselective NO synthase inhibitor N^ω^-nitro-L-arginine methylester (L-NAME), 10 μM of nonselective cyclooxygenase1/2 inhibitors (indomethacin), 1 mM of reactive oxygen species inhibitor (Tiron), 100 μM of selective NOX2 inhibitor (Apocynin), 5 μM of selective NOX4 inhibitor (GKT137831), and 10 μM of selective NOX2/4 inhibitor (VAS2870).

### Chemicals

All the drugs were purchased from Sigma (St. Louis, MO, USA).

### Statistical Analysis

Statistical analysis was performed by OriginPro version 8.0 for Windows (Origin Lab, Northampton, MA, USA). Data were presented as original recordings and a bar graph of the mean ± standard error of the mean (SEM); n indicates the number of animals or vessels being studied. Statistical comparison was performed with analysis of variance or repeated measures analysis of variance (ANOVA) followed by the Bonferroni post-hoc test. Half-maximal effective concentrations (*EC*_*50*_) of the response curve were fitted to sigmoidal functions. *E*_*max*_ is the maximal response to the agonist standardizes with 80 mM K^+^. The significance level was set at 0.05 in all tests.

## RESULTS

After 5 weeks, the body weights of FR groups were significantly lower compared to the control groups (FR40, 292.49±2.31 g; FR20, 344.51±4.08 g; control, 430.04±3.46 g; *p* < 0.001) **(Table 1)**. The organ weight of the FR groups was also significantly lower than the control groups **(Table 1)**. Mean arterial pressure (MAP) was significantly lower in FR40 groups as compared to the FR20 and control groups (FR40, 93.87±2.10 mmHg; FR20, 97.18±1.30 mmHg; control, 103.91±1.34 mmHg, *p* < 0.001) **(Figure 1C)**. The heart rate of FR40 groups was also significantly lower as compared to the FR20 and control groups (FR40, 304.75±12.46 beats; FR20, 318.63±5.24 beats; control, 363.05±11.88 beats; *p* < 0.01) **(Figure 1D)**.

**Table 1.**
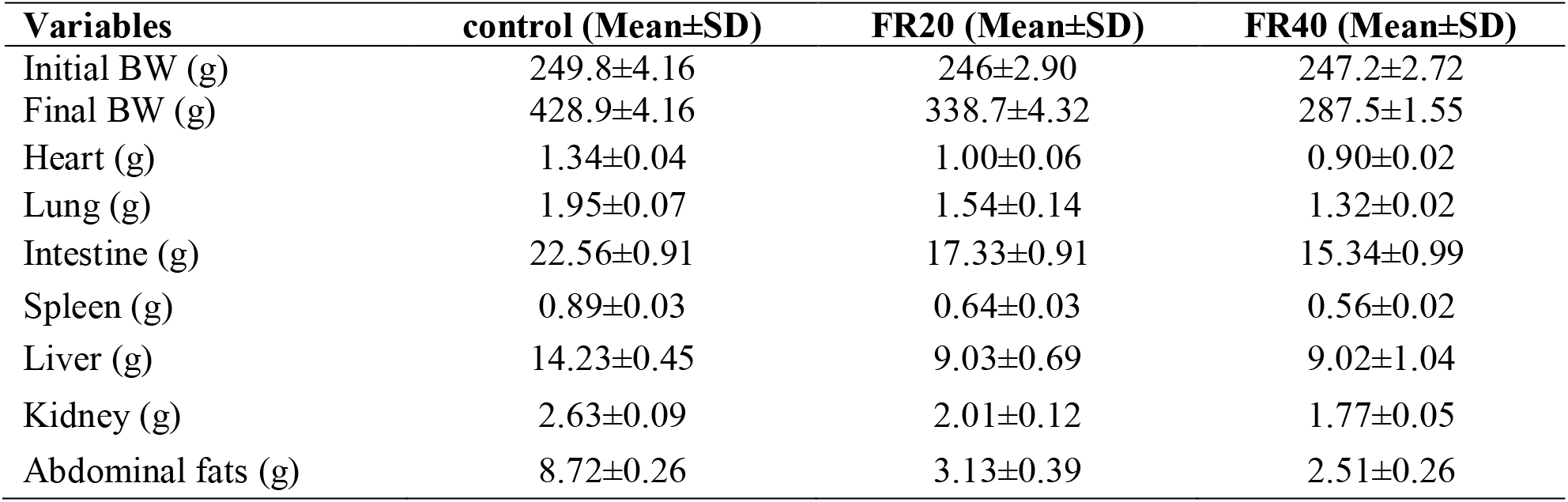
Body weight and proportional organ weights of all the groups after 5 weeks of food restriction

| Variables | control (Mean±SD) | FR20 (Mean±SD) | FR40 (Mean±SD) |
| --- | --- | --- | --- |
| Initial BW (g) | 249.8±4.16 | 246±2.90 | 247.2±2.72 |
| Final BW (g) | 428.9±4.16 | 338.7±4.32 | 287.5±1.55 |
| Heart (g) | 1.34±0.04 | 1.00±0.06 | 0.90±0.02 |
| Lung (g) | 1.95±0.07 | 1.54±0.14 | 1.32±0.02 |
| Intestine (g) | 22.56±0.91 | 17.33±0.91 | 15.34±0.99 |
| Spleen (g) | 0.89±0.03 | 0.64±0.03 | 0.56±0.02 |
| Liver (g) | 14.23±0.45 | 9.03±0.69 | 9.02±1.04 |
| Kidney (g) | 2.63±0.09 | 2.01±0.12 | 1.77±0.05 |
| Abdominal fats (g) | 8.72±0.26 | 3.13±0.39 | 2.51±0.26 |

**Figure 1.**
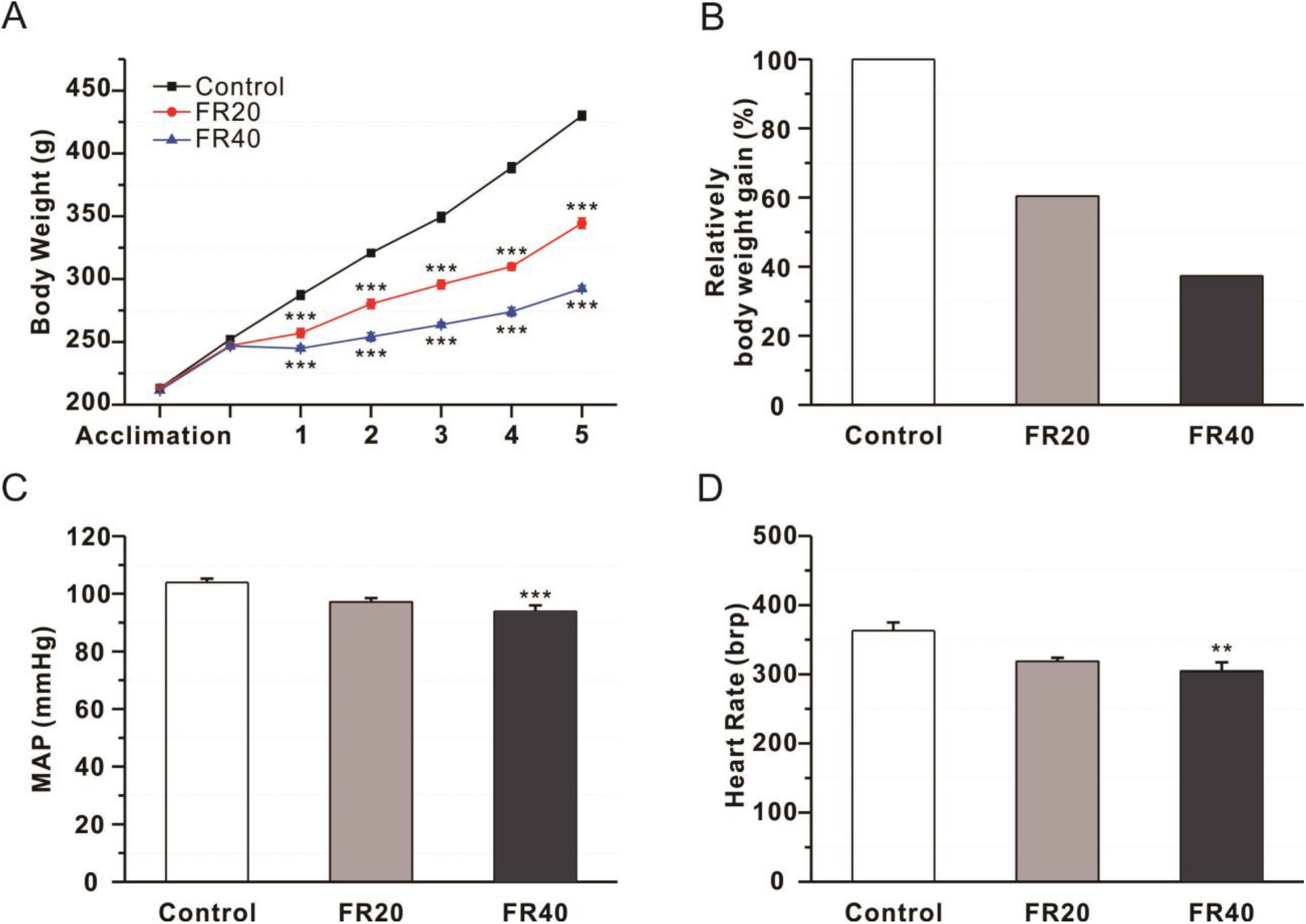
Summary of the body weight and blood pressure of the food restriction models. Monitoring was undertaken for 5 weeks of FR. Relative bodyweight gains were compared to the control (A, B). Mean arterial pressures (C) and heart rates (D) of the control, FR20, and FR40 of rats were also measured. Data is presented as mean±SEM and significant levels are indicated with \**p* < 0.05, \*\**p* < 0.01, and \*\*\**p* < 0.001.

### FR on Pulmonary Arterial Pressure

To understand the effect of FR on blood flow in the pulmonary circulation, we measured pulmonary arterial pressure via isolated perfusion lung experiments. FR did not affect the basal tone of PAP (FR40, 16.33±4.76 mmHg; FR20, 10.21±0.77 mmHg; control, 12.83±0.48 mmHg; *p* > 0.05) **(Figure 2A)**. Vasocontraction in the pulmonary artery by Ang II (FR40, 5.56±0.96 mmHg; FR20, 5.11±1.11 mmHg; control, 4.78±0.41 mmHg; *p* > 0.05) **(Figure 2B)**, hypoxia (FR40, 5.44±2.34 mmHg; FR20, 4.11±0.35 mmHg; control, 5.20±0.88 mmHg; *p* > 0.05) **(Figure 2C)** and high K^+^ (FR40, 38.79±5.09 mmHg; FR20, 33.03±5.23 mmHg; control, 48.92±3.86 mmHg; *p* > 0.05) **(Figure 2D)** exhibited no differences between the groups.

**Figure 2.**
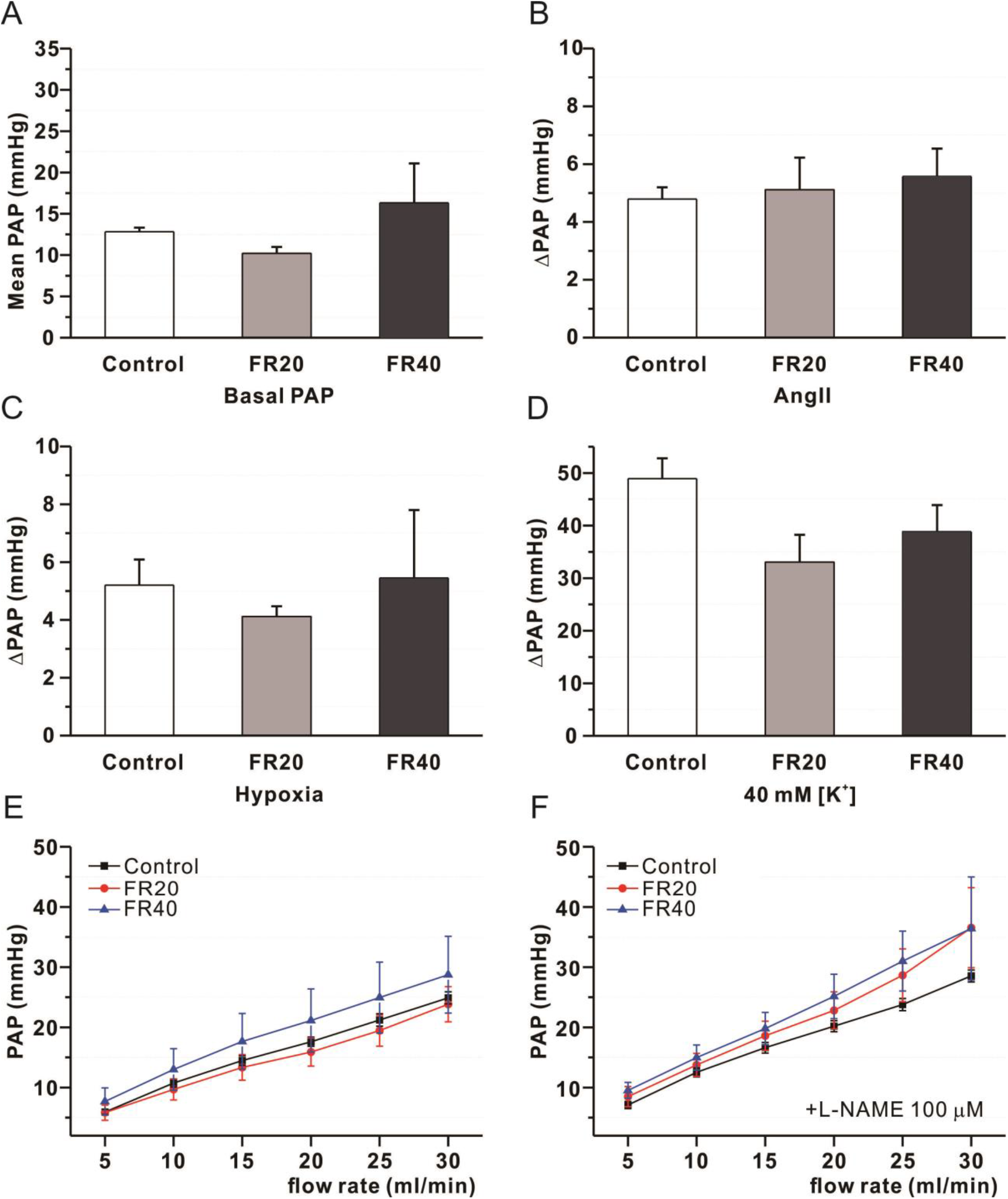
Effect of FR on the basal tone of pulmonary arterial pressure (PAP) Effect of FR on the basal tone of pulmonary arterial pressure (A), contractile response to Ang II at 1 *μ*g (B), hypoxia (O_2_ 3%) on hypoxia-induced pulmonary vasoconstriction (C) and on the contractile response to high K^+^-induced membrane depolarization on pulmonary arterial pressure (D) as compare between control, FR20, and FR40 groups. Flow rate is with, or without, inhibition of NOS (E, F). Data is presented as mean±SEM and significant levels are indicated with \**p* < 0.05, \*\**p* < 0.01, and \*\*\**p* < 0.001.

### Effect of FR on Vascular Smooth Muscle Reactivity

To understand the α1-adrenergic receptors in mesenteric arteries after 5 weeks of FR, we applied a dose-dependent response to PhE-mediated vasoconstriction in 5 minute intervals. We observed that short-term FR altered vascular contractility in mesenteric arteries. This alteration enhanced the sensitivity (*EC*_*50*_) to PhE-mediated vasoconstriction as compared to control (FR40, 1.09±0.10 μM; FR20, 1.39±0.07 μM; and control, 1.88±0.06 μM; *p <* 0.001) **(Figure 3F)**. The maximal constriction (*E*_*max*_) was significantly higher in the FR groups compared to control (FR40, 114.83±1.62%; FR20, 112.39±2.34%; control, 105.30±2.13%; *p <* 0.01) **(Figure 3D)**.

**Figure 3.**
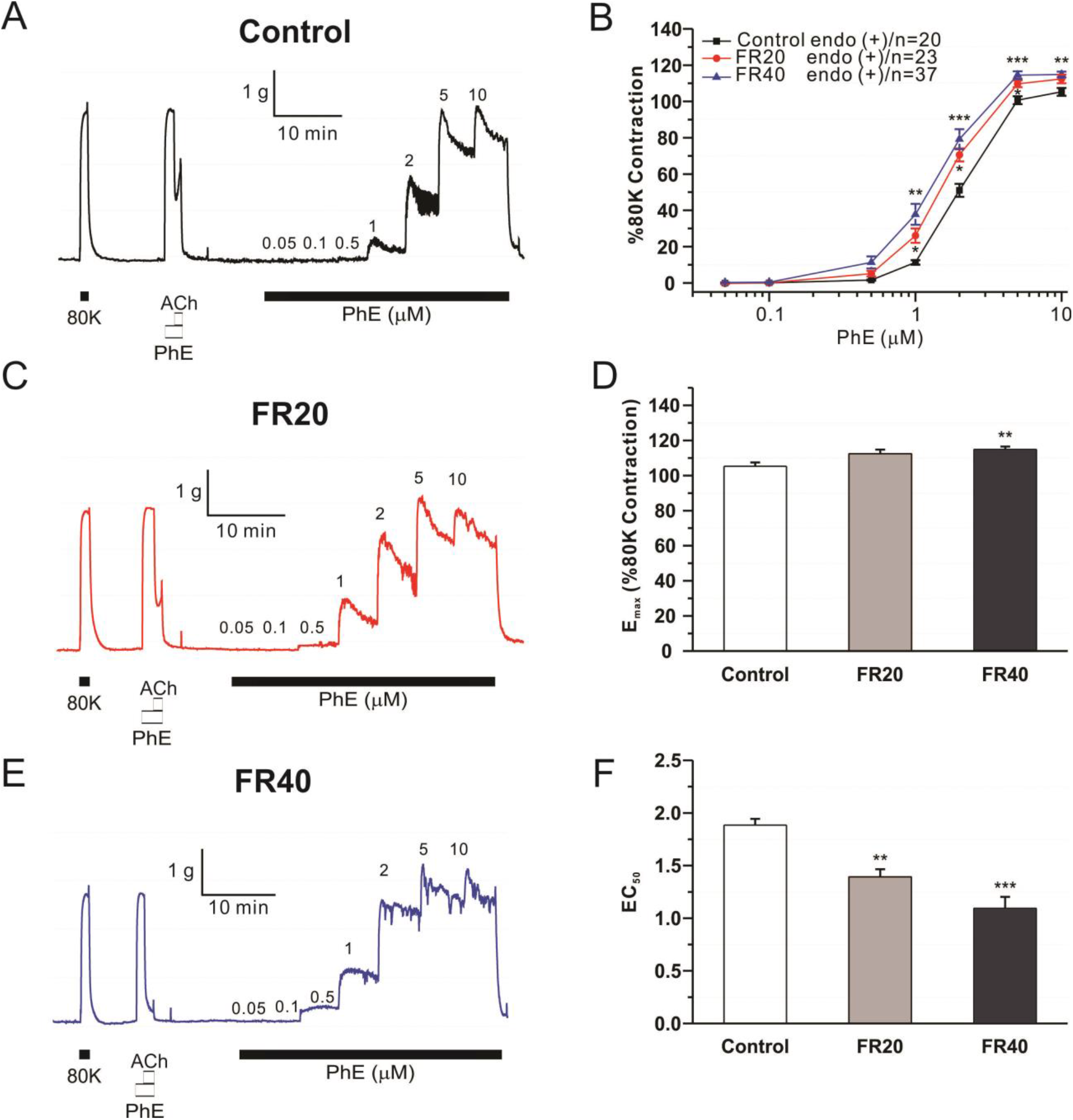
Effect of FR on PhE-mediated contraction in mesenteric arteries. Raw trace of PhE-mediated vasoconstriction in control (A), FR20 (C), and FR40 groups (E). Summary concentration-response curves for PhE-mediated vasoconstriction in the mesenteric artery (B). Mean values of maximal response (*E*_max_) and log *EC*_*50*_ derived from the experiment are shown in (D, F). The data is expressed as the percentage of the contractile response to 80 mM K^+^. Data is presented as mean±SEM and significant levels are indicated with \**p* < 0.05, \*\**p* < 0.01, and \*\*\**p* < 0.001.

We then tested whether FR influenced other vasoconstrictors such as serotonin (5-HT), thromboxane A2 (U46619), and high K^+^-induced vasoconstriction. FR had no effect on the sensitivity and maximal constriction to 5-HT **(Figure 4A, B)** and U46619 **(Figure 4C, D)**. Dose-dependent high K^+^ concentration was applied in 5 minute intervals from 20 to 120 mM. There were no differences in the sensitivity and maximal constriction between the groups **(Figure 4E, F)**.

**Figure 4.**
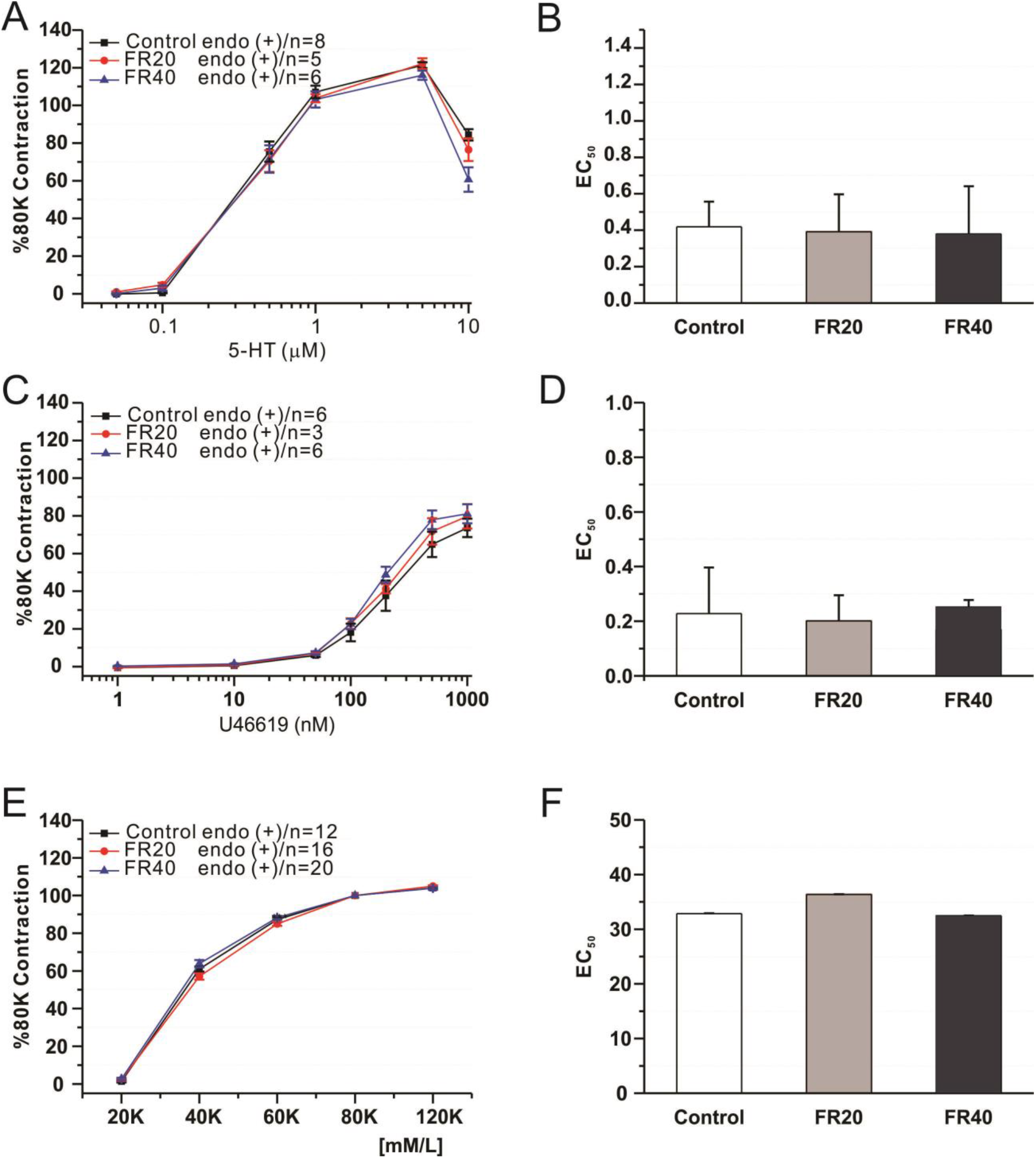
Effect of FR on contractile function of mesenteric smooth muscle cells. Effect of food restriction on dose-response curves and sensitivity to serotonin **(A, B)**, U46619 **(C, D)** and high K^+^ **(E, F)** in isolated mesenteric arteries of the groups. The data is expressed as the percentage of the contractile response to 80 mM [K^+^]. Data is presented as mean±SEM and significant levels are indicated with \**p* < 0.05, \*\**p* < 0.01, and \*\*\**p* < 0.001.

### Effect of FR on Endothelium-Dependent Relaxation

To determine the effect of FR on endothelial function, dose-dependent relaxation of ACh was performed. FR attenuated the ACh-induced vasorelaxation. The *EC*_*50*_ of FR groups was significantly higher compared to control (FR40, 0.37±0.03 μM; FR20, 0.31±0.03 μM; and control, 0.28±0.03 μM; *p <* 0.05) **(Figure 5D)**. However, there was no difference in maximal relaxation between the groups (FR40, 84.24±2.35%; FR20, 87.64±2.22%; and control, 89.04±1.71%; *p >* 0.05) **(Figure 5F)**.

**Figure 5.**
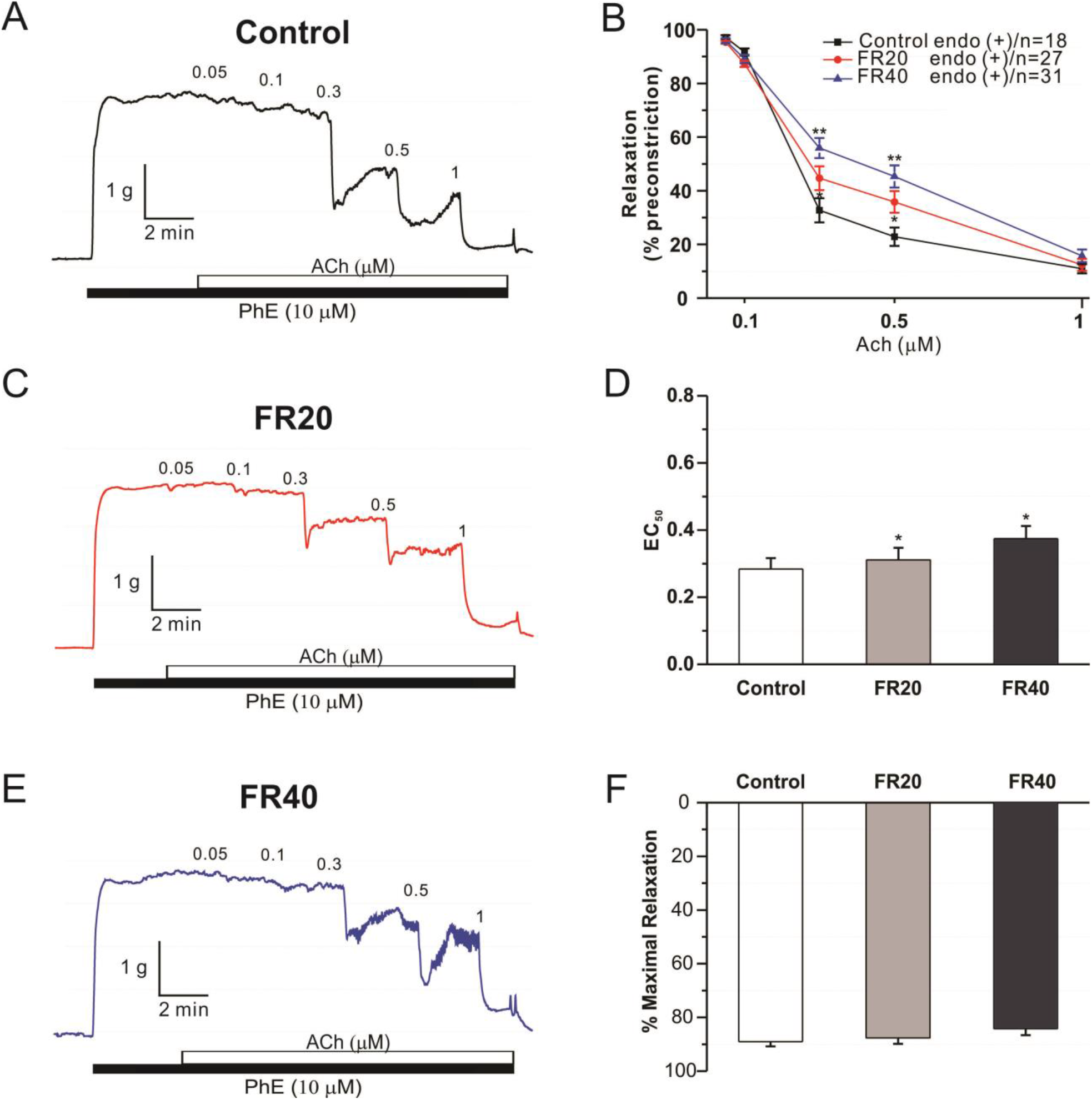
Effect of FR on endothelium-dependent relaxation in mesenteric arteries. Effect of FR on endothelium-dependent relaxation from control (A), FR20 (C), and FR40 (E) isolated mesenteric arteries. The data is expressed as the percentage of relaxation relative to the precontracted state with PhE. Summary concentration-response curves for ACh-induced vasodilation in the mesenteric artery (B). Mean values of maximal response (*E*_max_) and log *EC*_*50*_ derived from the experiment shown in (D, F). Data is presented as mean±SEM and significant levels are indicated with \**p* < 0.05, \*\**p* < 0.01, and \*\*\**p* < 0.001.

### Effect of FR on NOS and Endothelium-Independent Relaxation

We assessed the basal NOS activity by 100 μM of L-NAME with PhE pre-contraction to ∼15% of 80K. Without the pre-contractor, L-NAME produced no response in the isolated mesenteric artery **(Figure 6B)**. There were no differences between the groups **(Figure 6C)**, which indicated that FR induced endothelial dysfunction was not due to alteration of basal NOS activity. Endothelium-independent relaxation by sodium nitroprusside (SNP) showed a similar relaxation between the groups **(Figure 6D)**.

**Figure 6.**
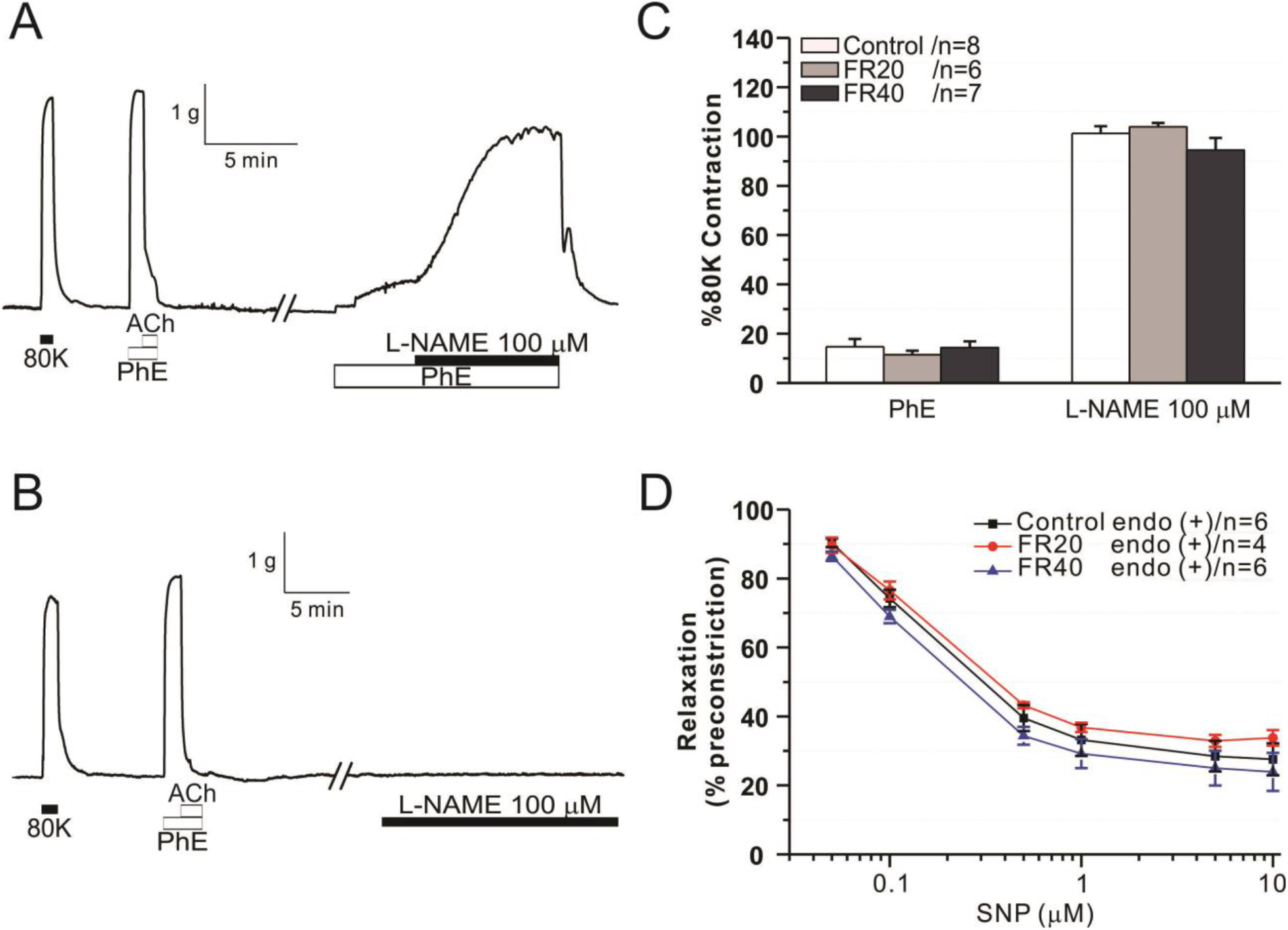
Effect of FR on basal NOS and endothelium-independent relaxation in mesenteric arteries. Effect of FR on NOS activity of isolated mesenteric arteries. This shows the response of L-NAME with PhE pretone in the mesenteric artery (A). There was no response of L-NAME without PhE pretone in the mesenteric artery (B). The data is expressed as a percentage of the contractile response to 80 mM K^+^ (C). Endothelium-independent relaxation of isolated mesenteric arteries. The data are expressed as the percentage of relaxation relative to pre-contracted with PhE. Summary concentration-response curves for SNP-induced vasodilation in the mesenteric artery (D). Data is presented as mean±SEM and significant levels are indicated with \**p* < 0.05, \*\**p* < 0.01, and \*\*\**p* < 0.001.

### Effect of FR on eNOS Mediated Vasorelaxation

To clarify the understanding of the role of eNOS with regards to endothelium-dependent relaxation in mesenteric arteries, we performed a dose-response with ACh in the presence of 10 μM of indomethacin (COX inhibitor) to inhibit the production of prostacyclin with a small and intermediate conductance Ca^2+^ activated K^+^ channel blockers (100 μM of apamin and 1 μM of TRAM-34). In the presence of indomethacin+apamin+TRAM-34 for 30 minutes, it was shown that vasorelaxation was suppressed in each group **(Figure 7A, C, E)**. However, an eNOS-dependent response to ACh in the mesenteric arteries of FR groups remains impaired, in terms of sensitivity and maximal response when compared to the control groups **(Figure 7D, F)**.

**Figure 7.**
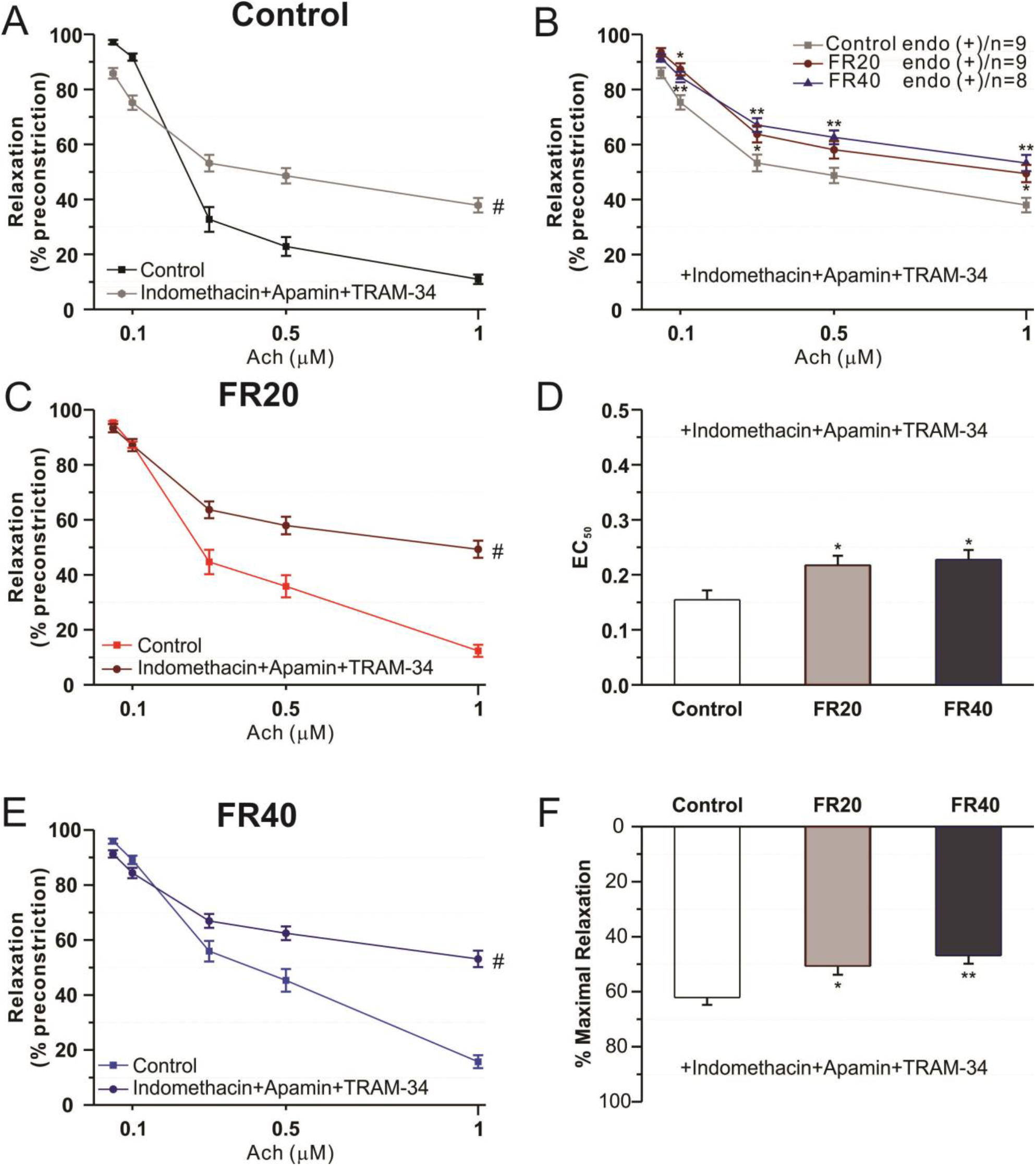
Effect of FR on eNOS-dependent vasorelaxation in mesenteric arteries. Effect of FR on eNOS-dependent vasorelaxation. This shows summary concentration-response curves to ACh in the absence (control) and presence of indomethacin (10 μM) +apamin (100 μM) +TRAM-34 (1 μM) of control (A), FR20 (C) and FR40 (E). Dose-response curves and sensitivity to ACh in the presence of indomethacin+apamin+TRAM-34 (B, D, F). Data is presented as mean±SEM and significant levels are indicated with \**p* < 0.05, \*\**p* < 0.01, and \*\*\**p* < 0.001.

### Effect of Non-Specific COX and NOS Inhibitors on PhE-Mediated Vasoconstriction

To investigate the role of prostanoids in FR enhanced vascular contractility, 10 μM of indomethacin was introduced before application of PhE. FR groups displayed greater maximal constriction and higher sensitivity when compared to the control groups **(Figure 8A, B)**.

**Figure 8.**
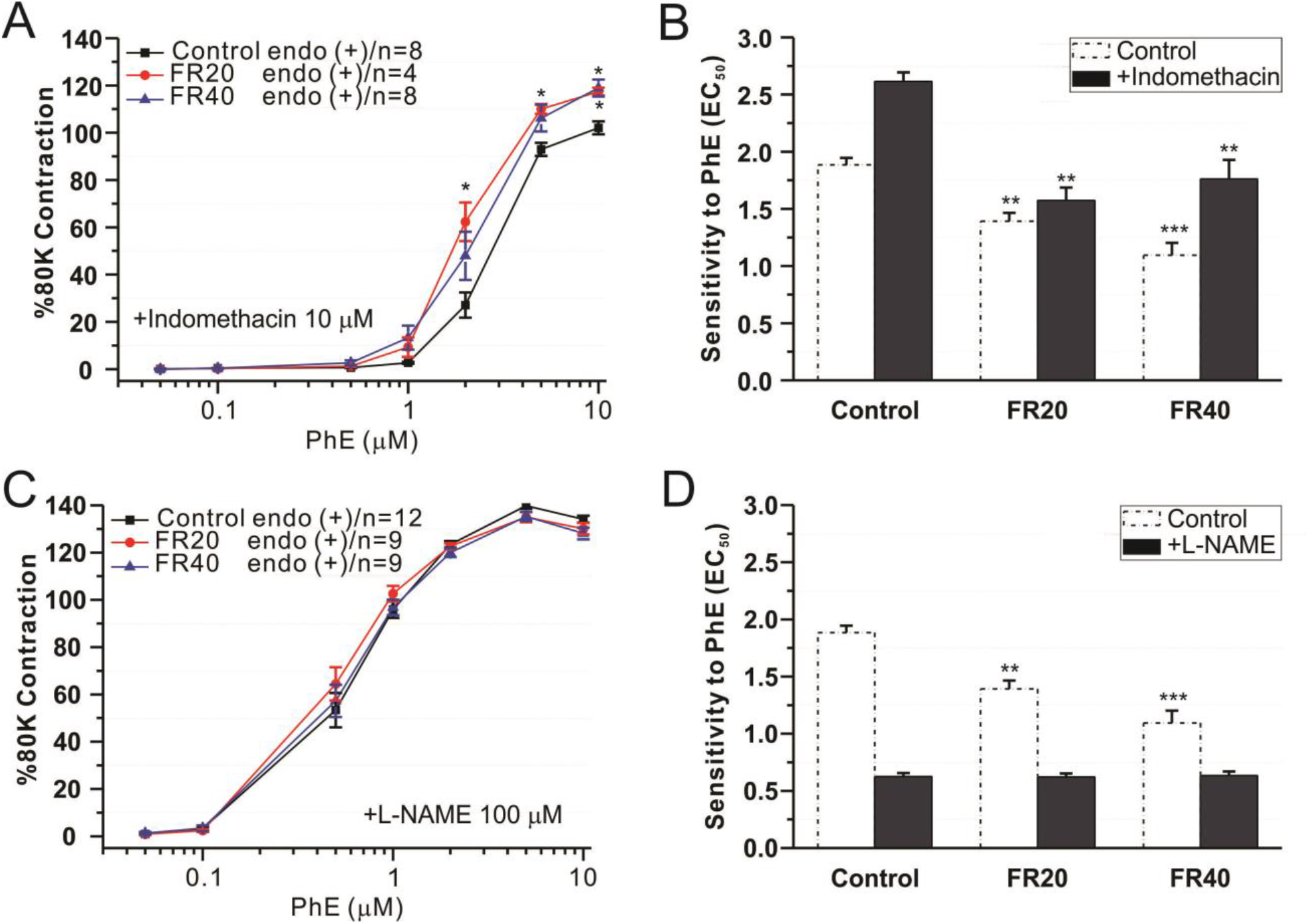
Effect of non-specific COX and NOS inhibitors in mesenteric arteries. Effect of FR on PhE-induced contractile response in the presence of indomethacin (10 μM) and NOS inhibitor (100 μM). Summary concentration-response curves to PhE in the presence of indomethacin (A, B) and L-NAME (C, D). The data is expressed as the percentage of the contractile response to 80 mM K^+^. Data is presented as mean±SEM and significant levels are indicated with \**p* < 0.05, \*\**p* < 0.01, and \*\*\**p* < 0.001.

To understand whether eNOS might contribute to the enhancement of the PhE-mediated vasoconstriction in mesenteric arteries, a non-specific NOS inhibitor were applied (100 μM of L-NAME for 30 minutes) before administering dose-dependent of PhE. Inhibition of NOS increased contractility of PhE in each groups and inhibited further constriction with regards to FR groups **(Figure 8C, D)**.

### Effect of Specific NOXs and ROS Scavenger Inhibitors on PhE-mediated vasoconstriction

To investigate the role of NOX in the enhancement of PhE-mediated vasoconstriction on the mesenteric arteries in FR groups, arteries were incubated with specific NOXs inhibitors for 30 minutes prior to the application of PhE. NOXs are a major source of ROS production in the vascular system. Among the seven NOX isoforms, NOX 1, 2, 4 and 5 are expressed in vascular endothelial cells. In the presence of NOX2 inhibitor (100 μM of apocynin), NOX4 inhibitor (5 μM of GKT137831), and NOX2/4 (10 μM of VAS2870), PhE further suppressed constriction in FR groups. There were no statistically significant differences found in sensitivity and the maximal constriction in the presence of NOXs inhibitor **(Figure 9B, D, and F)**.

**Figure 9.**
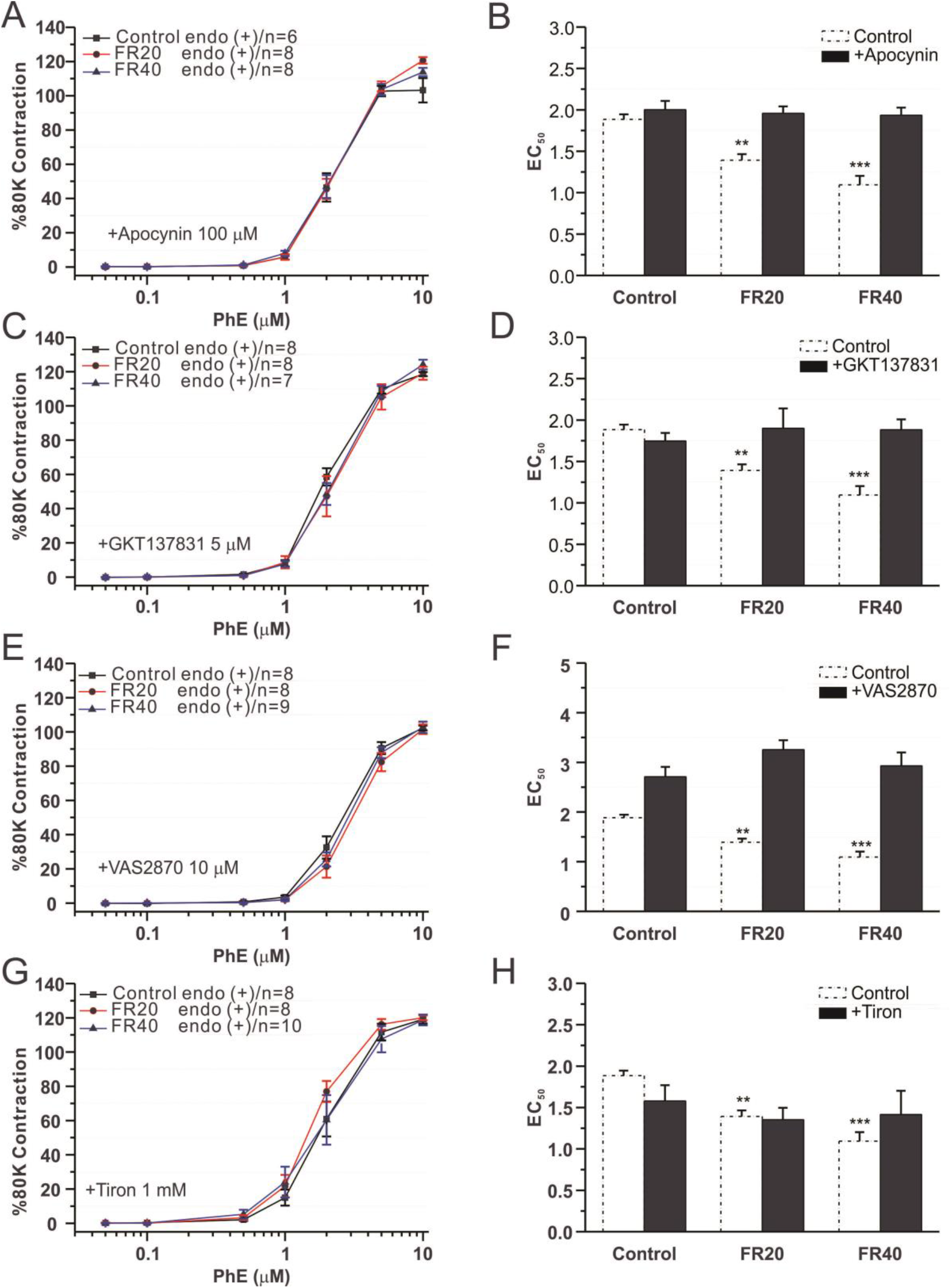
Effect of specific NOXs and ROS inhibitors in mesenteric arteries. Effect of FR on PhE-induced contractile response in the presence of NOXs inhibitors [apocynin (100 μM), GKT137831 (5 μM), and VAS2870 (10 μM)] and ROS inhibitor (Tiron, 1 mM). Summary concentration-response curves and sensitivity to PhE in the presence of apocynin (A, B); GKT137831 (C, D); VAS2870 (E, F); and Tiron (G, H). The data is expressed as the percentage of the contractile response to 80 mM K^+^. Data is presented as mean±SEM and significant levels are indicated with \**p* < 0.05, \*\**p* < 0.01, and \*\*\**p* < 0.001.

To determine whether the generation of ROS played a role in mediating the effect of PhE-induced higher constriction in FR groups, a ROS scavenger (Tiron, 1 mM) was introduced 30 minutes prior to the application of dose-response curves of PhE. After the ROS scavenger was introduced, PhE further suppressed constriction in FR groups. There were no statistically significant differences in sensitivity and maximal constriction **(Figure 9G, H)**.

## DISCUSSION

The main findings of this study are that FR is associated with bradycardia and hypotension; the enhancement α1-adrenergic mediated contractility, the impairment of eNOS mediated endothelium-dependent relaxation; and the alteration of NOXs/ROS activity in mesenteric arteries.

FR influenced body and organ weight in the heart, liver, lung, spleen, kidneys and intestine. In contrast, the brain, a vital organ, showed no differences between the groups. This could be a pattern of circulatory redistribution aimed at preservation of brain growth. FR reduced the body weight by 19% in FR20 and 62.6% in FR40 compared to control groups. A previous study suggested that caloric restriction reduced cardiac hypertrophy in the hypertension rat model [18]. In ventilated and perfused lungs in this study, PAP and HPV did not differ response to FR. HPV is a physiological compensatory phenomenon by the pulmonary artery to prevent arterial hypoxemia by diverting the blood flow from poorly ventilated regions to normoxic alveoli [33, 34]. Our findings showed that FR did not have an effect on mean pulmonary arterial pressure. In addition, there were no statistically significant differences in response to various vasoconstrictors, which is consistent with a previous study in the control model [18].

Severe FR induced a lower heart rate and hypotension in female rats [26], which is consistent with our finding in the male FR model. We found that FR enhanced vascular contractility in response to α1-adrenoreceptor agonist despite observed hypotension. We propose that the alteration of receptor density may not be the primary mechanism, but is counterbalanced by the cardiovascular system maintenance of blood flows in the bodies responses to FR induced hypotension. Another plausible explanation is that in order to maintain blood circulation in the body, the mesenteric arteries increase their vascular tone to prevent organs from being damaged by hypovolemic shocks. It is well known that increases in adrenergic agonists, such as α1−adrenergic receptors, regulates vasoconstriction in VSMCs in order to maintain blood flow and thus plays a significant contributory role in the cardiovascular system [35]. Our findings were consistent with recent studies in female rats [24, 26].

A previous study has shown that plasma nitric oxide was reduced in FR groups [24]. This could be explained by the attenuation of endothelium-dependent relaxation in our study. The endothelium plays a significant role in regulating vascular tone by releasing endothelium-derived NO [6], prostacyclin [36], and endothelium-dependent hyperpolarization factors [37]. Reductions in the production of NO from endothelium impaired vasodilation and augmented the vasoconstriction which is the common pathway for CVD [6]. The eNOS is the main pathway involved in endothelium-dependent relaxation in response to ACh [38]. To understand our hypothesis that FR altered eNOS function in the mesenteric artery, we inhibited the other possible pathways. After, blockage with indomethcin+apamin+TRAM-34 for 30 minutes, the ACh remained significantly lower compared to the control group. We suggest that FR is associated with alteration of the eNOS-NO pathway in mesenteric arteries, which results in reduced plasma NO production [24]. We investigated the eNOS contribution to α1-adrenergic mediated contraction by introducing a nonspecific NOS inhibitor before the application of α1-adrenergic contraction. In the presence of NOS inhibitor, there was no longer a significant difference between the groups. FR might reduce the eNOS function to increase vascular smooth muscle contraction for maintaining the blood flow.

We hypothesized that the alteration of eNOS function might be related to the enhancement of NOXs activity in vascular endothelial cells, which inactivated NO. Increased ROS production, leading to mitochondrial dysfunction have been reported in AN women [39]. Out of seven NOXs, isoform, NOX2/4 may responsible for the enhancement of vascular contractility [13]. We investigated the role of NOX2/4 in α1-adrenergic mediated contraction with apocynin, GKT137831, VAS2870, and Tiron for 30 minutes. In the presence of NOXs and ROS inhibitors, the differences of α1-adrenergic mediated contraction in isolated mesenteric arteries were eliminated. Our results suggest that FR might enhance superoxide anion production from NOX2/4, which leads to augmentation of vascular contraction. A similar finding was reported in 50% of FR in female offspring [40].

### Limitation and physiological implication of the study

According to our study, we found an interesting physiological effect of the cardiovascular system response to severe food restriction in males. Augmentation of α1-adrenoreceptor agonist might be a counterbalance by the mesenteric artery to prevent organs from being damaged due to lower blood pressure. Notably, in order to increase the vascular tone, endothelium might suppress the eNOS enzyme production which leads to decreased vasodilation agents such as nitric oxide. A limitation of our study was that we did not check α1-adrenoreceptor agonist in pulmonary vascular function. A previous study suggested that there was an increase in α1-adrenoreceptor agonist in female responses to FR [24].

### Conclusion

The effect of FR on pulmonary and mesenteric vascular function requires further investigation. FR was associated with lower mean arterial pressure, but not pulmonary arterial pressure. Interestingly, FR may suppress eNOS function by enhancing NOXs activity in mesenteric arteries by stimulating α1-adrenergic mediated vasoconstriction in mesenteric arteries. This could be a vascular physiology compensation in response to FR.

## DATA AVAILABILITY STATEMENT

The datasets generated for this study are available on request to the corresponding author.

## ETHICAL STATEMENT

The animal studies were performed in accordance with the National Institutes of Health guidelines for the care and use of animals, and the Institutional Animal Care and Use Committee (IACUC) in Chung-Ang University (IACUC approval no. 2019-00047).

## AUTHOR CONTRIBUTIONS

RV designed and conceptualized the study, performed the experiment and data analysis and contributed to the manuscript intellectual content. HY designed and conceptualized the study; interpreted the data; and revised the manuscript intellectual content.

## ACKNOWLEDGMENT

This study was part of a PhD dissertation which was submitted to Chung-Ang University Library. Part of the abstract was published as an abstract in the Circulation journal. We thank the NIH Library Writing Center for manuscript editing assistance.

